# Sequence capture of ultraconserved elements from bird museum specimens

**DOI:** 10.1101/020271

**Authors:** John E. McCormack, Whitney L.E. Tsai, Brant C. Faircloth

## Abstract

New DNA sequencing technologies are allowing researchers to explore the genomes of the millions of natural history specimens collected prior to the molecular era. Yet, we know little about how well specific next-generation sequencing (NGS) techniques work with the degraded DNA typically extracted from museum specimens. Here, we use one type of NGS approach, sequence capture of ultraconserved elements (UCEs), to collect data from bird museum specimens as old as 120 years. We targeted approximately 5,000 UCE loci in 27 Western Scrub-Jays (*Aphelocoma californica*) representing three evolutionary lineages, and we collected an average of 3,749 UCE loci containing 4,460 single nucleotide polymorphisms (SNPs). Despite older specimens producing fewer and shorter loci in general, we collected thousands of markers from even the oldest specimens. More sequencing reads per individual helped to boost the number of UCE loci we recovered from older specimens, but more sequencing was not as successful at increasing the length of loci. We detected contamination in some samples and determined contamination was more prevalent in older samples that were subject to less sequencing. For the phylogeny generated from concatenated UCE loci, contamination led to incorrect placement of some individuals. In contrast, a species tree constructed from SNPs called within UCE loci correctly placed individuals into three monophyletic groups, perhaps because of the stricter analytical procedures we used for SNP calling. This study and other recent studies on the genomics of museums specimens have profound implications for natural history collections, where millions of older specimens should now be considered genomic resources.

## Introduction

Natural history collections house billions of specimens worldwide, providing a record of Earth’s biodiversity in space and time and a source of data to answer questions about evolution, ecology, conservation, and human health (Austin & Melville, 2006; Suarez & Tsutsui, 2004; Winker, 2004). Modern specimen preparation methods include archiving frozen tissue for DNA analysis, yet the millions of specimens collected prior to the molecular age lack this ready source of high quality DNA.

We have learned much during the last 25 years of studying ancient DNA from museum specimens using older DNA sequencing technology like Sanger sequencing (Cooper *et al.*, 1992; Houde & Braun, 1988; Mundy *et al.*, 1997; Thomas *et al.*, 1989; Thomas *et al.*, 1990; Wayne *et al.*, 1999), but it is also well documented that Sanger protocols are laborious and have many limitations (Soltis & Soltis, 1993; Wandeler *et al.*, 2007). Museum conditions are optimized for whole specimen longevity, not molecular stability, so DNA extracted directly from museum specimens is often fragmented and damaged in various ways (Axelsson *et al.*, 2008; Briggs *et al.*, 2007; Briggs *et al.*, 2010; Molak & Ho, 2011), and it also almost certainly contains contaminant DNA (Hofreiter *et al.*, 2001; Pääbo, 1989). For traditional Sanger sequencing, these short (and potentially contaminant) DNA fragments must be targeted and processed individually, greatly multiplying the workload needed to collect the same amount of genetic data from better preserved starting material.

Next-generation DNA sequencing (NGS) offers a more efficient way of sequencing DNA from museum specimens (Hofreiter *et al.*, 2015; Rizzi *et al.*, 2012). Samples can be sequenced in parallel, and current instruments generate many millions of reads per sample. First applied to sequence the mammoth genome (Poinar *et al.*, 2006), scientists have also used NGS to sequence the Neanderthal genome from bone fragments, taking advantage of the millions of sequence reads to overcome the problem of contamination from modern humans that would otherwise have reduced the overall number of Neanderthal DNA reads to an unacceptably low number (Green *et al.*, 2010; Meyer *et al.*, 2012). Other applications of NGS to museum specimens include research on the evolutionary history of species like primates, for which it would be difficult or unethical to collect specimens today (Guschanski *et al.*, 2013), and time series studies (Habel *et al.*, 2014) that examined how genetic diversity has changed in wild rodent populations with shifting elevational distributions caused by climate change (Bi *et al.*, 2013; Rowe *et al.*, 2011). New protocols and technological advances continue to improve the collection of whole genomes from ancient samples (Der Sarkissian *et al.*, 2015; Staats *et al.*, 2013).

Despite these exciting advances, there are many ways to use NGS in concert with ancient DNA, and it is not yet clear how DNA degradation might impair the collection of data using different approaches (Knapp & Hofreiter, 2010). Whole-genome shotgun sequencing is fairly straightforward, usually involving only a library preparation step of the already fragmented DNA (Besnard *et al.*, 2015; Besnard *et al.*, 2014; Zedane *et al.*, 2015). Protocols targeting subsets of the genome are more complicated because they involve more steps to isolate the targeted subset of loci from the rest of the genome. For example, extensive age-related DNA fragmentation might negatively affect restriction-associated digest sequencing (RADseq) or genotyping by sequencing (GBS) approaches, because this family of techniques relies on making one or two systematic cuts to genomic DNA in precise locations (Baird *et al.*, 2008; Etter *et al.*, 2011). Similarly, DNA fragmentation might limit the utility of target enrichment techniques (also known as sequence capture; Gnirke *et al.*, 2009; Mamanova *et al.*, 2010) if the sequenced reads are too short to capture or captured reads are too short to reassemble into longer loci. Capture approaches have proven effective for analyses of high-copy ancient mitochondrial DNA from museum specimens (Horn, 2012; Mason *et al.*, 2011; Thalmann *et al.*, 2013; Vilstrup *et al.*, 2013), but low-copy nuclear DNA is likely to pose more problems and is less tested.

Our goal was to assess the effectiveness of a specific kind of NGS approach, sequence capture of ultraconserved elements (UCEs), for collecting genomic data from museum specimens for phylogenetics and population genetics. UCEs are a class of highly conserved and abundant nuclear marker distributed throughout the genomes of most organisms (Bejerano *et al.*, 2004; Reneker *et al.*, 2012; Siepel *et al.*, 2005; Stephen *et al.*, 2008). UCEs work because synthetic, oligonucleotide probes bind to the central, conserved portion of each UCE locus, allowing researchers to isolate and sequence this region as well as more variable, flanking DNA that is also captured (Faircloth *et al.*, 2012). Sequence variation in the core region is useful for deep phylogenetic questions (Crawford *et al.*, 2012; McCormack *et al.*, 2013), whereas variation in the flanking regions is useful at phylogeographic or population genetic timescales (Smith *et al.*, 2014). One useful feature of UCEs is that they are found across a variety of taxa including amniotes, fish (Faircloth *et al.*, 2013), and insects (Faircloth *et al.*, 2014), making the development of new probe sets for many groups unnecessary.

Although the UCE approach is theoretically possible with ancient DNA, the outcome may be adversely affected by age-related DNA degradation. For example, the UCE protocol involves capturing small (60-120 base pair), conserved genomic targets which could be rendered too small to capture by age-related fragmentation. Additionally, sequence variation sufficient to resolve moderate and shallow-level relationships is concentrated in the flanking regions of UCE loci, which might be impossible to assemble from short, degraded DNA fragments. On the other hand, if the UCE capture approach can be effectively applied to museum specimens, then this technique, with a pre-existing probe set that routinely captures many thousands of loci across vertebrates, would immediately open the door to collecting genome-scale data from the millions of specimens housed in museums.

## Methods

### Study species, sampling, and DNA extraction

We cut toe pads from 27 study skins (Table 1) spanning three divergent evolutionary lineages of the Western Scrub-Jay (*Aphelocoma californica*). One lineage is found along the Pacific coast (*californica* group), one in the interior United States and northern Mexico (*woodhouseii* group), and one in southern Mexico (*sumichrasti* group). These three lineages are divergent in both DNA and appearance (Delaney *et al.*, 2008; Gowen *et al.*, 2014; Peterson, 1992) and diverged from one another within the last 0.5 to 2.5 million years (McCormack *et al.*, 2011).

**Table 1.**
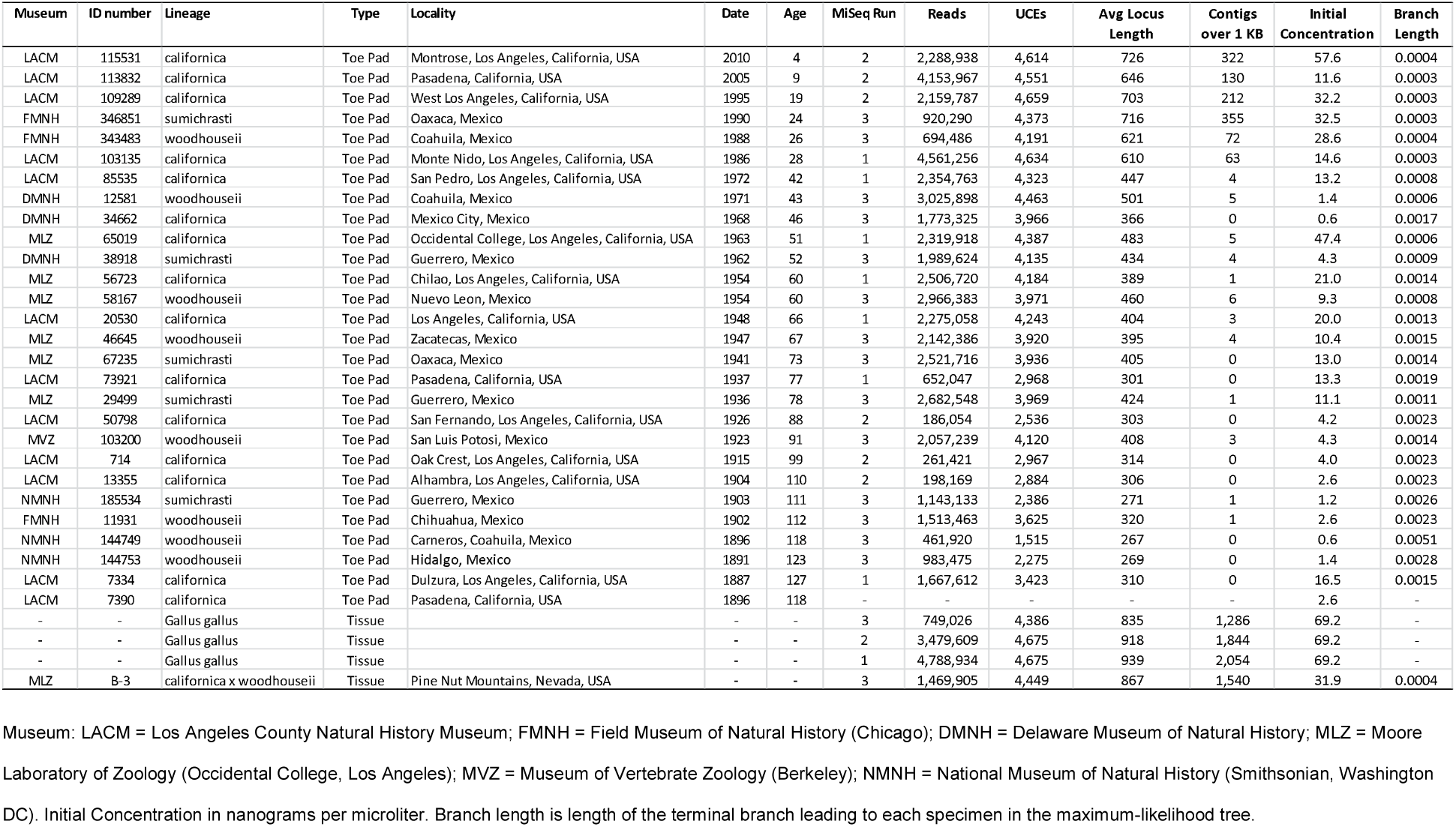
Information on specimens, sequencing, and results.

| Museum | ID number | Lineage | Type | Locality | Date | Age | MiSeq Run | Reads | UCEs | Avg Locus Length | Contigs over 1 KB | Initial Concentration | Branch Length |
| --- | --- | --- | --- | --- | --- | --- | --- | --- | --- | --- | --- | --- | --- |
| LACM | 115531 | californica | Toe Pad | Montrose, Los Angeles, California, USA | 2010 | 4 | 2 | 2,288,938 | 4,614 | 726 | 322 | 57.6 | 0.0004 |
| LACM | 113832 | californica | Toe Pad | Pasadena, California, USA | 2005 | 9 | 2 | 4,153,967 | 4,551 | 646 | 130 | 11.6 | 0.0003 |
| LACM | 109289 | californica | Toe Pad | West Los Angeles, California, USA | 1995 | 19 | 2 | 2,159,787 | 4,659 | 703 | 212 | 32.2 | 0.0003 |
| FMNH | 346851 | sumichrasti | Toe Pad | Oaxaca, Mexico | 1990 | 24 | 3 | 920,290 | 4,373 | 716 | 355 | 32.5 | 0.0003 |
| FMNH | 343483 | woodhouseii | Toe Pad | Coahuila, Mexico | 1988 | 26 | 3 | 694,486 | 4,191 | 621 | 72 | 28.6 | 0.0004 |
| LACM | 103135 | californica | Toe Pad | Monte Nido, Los Angeles, California, USA | 1986 | 28 | 1 | 4,561,256 | 4,634 | 610 | 63 | 14.6 | 0.0003 |
| LACM | 85535 | californica | Toe Pad | San Pedro, Los Angeles, California, USA | 1972 | 42 | 1 | 2,354,763 | 4,323 | 447 | 4 | 13.2 | 0.0008 |
| DMNH | 12581 | woodhouseii | Toe Pad | Coahuila, Mexico | 1971 | 43 | 3 | 3,025,898 | 4,463 | 501 | 5 | 1.4 | 0.0006 |
| DMNH | 34662 | californica | Toe Pad | Mexico City, Mexico | 1968 | 46 | 3 | 1,773,325 | 3,966 | 366 | 0 | 0.6 | 0.0017 |
| MLZ | 65019 | californica | Toe Pad | Occidental College, Los Angeles, California, USA | 1963 | 51 | 1 | 2,319,918 | 4,387 | 483 | 5 | 47.4 | 0.0006 |
| DMNH | 38918 | sumichrasti | Toe Pad | Guerrero, Mexico | 1962 | 52 | 3 | 1,989,624 | 4,135 | 434 | 4 | 4.3 | 0.0009 |
| MLZ | 56723 | californica | Toe Pad | Chilao, Los Angeles, California, USA | 1954 | 60 | 1 | 2,506,720 | 4,184 | 389 | 1 | 21.0 | 0.0014 |
| MLZ | 58167 | woodhouseii | Toe Pad | Nuevo Leon, Mexico | 1954 | 60 | 3 | 2,966,383 | 3,971 | 460 | 6 | 9.3 | 0.0008 |
| LACM | 20530 | californica | Toe Pad | Los Angeles, California, USA | 1948 | 66 | 1 | 2,275,058 | 4,243 | 404 | 3 | 20.0 | 0.0013 |
| MLZ | 46645 | woodhouseii | Toe Pad | Zacatecas, Mexico | 1947 | 67 | 3 | 2,142,386 | 3,920 | 395 | 4 | 10.4 | 0.0015 |
| MLZ | 67235 | sumichrasti | Toe Pad | Oaxaca, Mexico | 1941 | 73 | 3 | 2,521,716 | 3,936 | 405 | 0 | 13.0 | 0.0014 |
| LACM | 73921 | californica | Toe Pad | Pasadena, California, USA | 1937 | 77 | 1 | 652,047 | 2,968 | 301 | 0 | 13.3 | 0.0019 |
| MLZ | 29499 | sumichrasti | Toe Pad | Guerrero, Mexico | 1936 | 78 | 3 | 2,682,548 | 3,969 | 424 | 1 | 11.1 | 0.0011 |
| LACM | 50798 | californica | Toe Pad | San Fernando, Los Angeles, California, USA | 1926 | 88 | 2 | 186,054 | 2,536 | 303 | 0 | 4.2 | 0.0023 |
| MVZ | 103200 | woodhouseii | Toe Pad | San Luis Potosi, Mexico | 1923 | 91 | 3 | 2,057,239 | 4,120 | 408 | 3 | 4.3 | 0.0014 |
| LACM | 714 | californica | Toe Pad | Oak Crest, Los Angeles, California, USA | 1915 | 99 | 2 | 261,421 | 2,967 | 314 | 0 | 4.0 | 0.0023 |
| LACM | 13355 | californica | Toe Pad | Alhambra, Los Angeles, California, USA | 1904 | 110 | 2 | 198,169 | 2,884 | 306 | 0 | 2.6 | 0.0023 |
| NMNH | 185534 | sumichrasti | Toe Pad | Guerrero, Mexico | 1903 | 111 | 3 | 1,143,133 | 2,386 | 271 | 1 | 1.2 | 0.0026 |
| FMNH | 11931 | woodhouseii | Toe Pad | Chihuahua, Mexico | 1902 | 112 | 3 | 1,513,463 | 3,625 | 320 | 1 | 2.6 | 0.0023 |
| NMNH | 144749 | woodhouseii | Toe Pad | Carneros, Coahuila, Mexico | 1896 | 118 | 3 | 461,920 | 1,515 | 267 | 0 | 0.6 | 0.0051 |
| NMNH | 144753 | woodhouseii | Toe Pad | Hidalgo, Mexico | 1891 | 123 | 3 | 983,475 | 2,275 | 269 | 0 | 1.4 | 0.0028 |
| LACM | 7334 | californica | Toe Pad | Dulzura, Los Angeles, California, USA | 1887 | 127 | 1 | 1,667,612 | 3,423 | 310 | 0 | 16.5 | 0.0015 |
| LACM | 7390 | californica | Toe Pad | Pasadena, California, USA | 1896 | 118 | - | - | - | - | - | 2.6 | - |
| - | - | Gallus gallus | Tissue |  | - | - | 3 | 749,026 | 4,386 | 835 | 1,286 | 69.2 | - |
| - | - | Gallus gallus | Tissue |  | - | - | 2 | 3,479,609 | 4,675 | 918 | 1,844 | 69.2 | - |
| - | - | Gallus gallus | Tissue |  | - | - | 1 | 4,788,934 | 4,675 | 939 | 2,054 | 69.2 | - |
| MLZ | B-3 | californica x woodhouseii | Tissue | Pine Nut Mountains, Nevada, USA | - | - | 3 | 1,469,905 | 4,449 | 867 | 1,540 | 31.9 | 0.0004 |

Museum: LACM = Los Angeles County Natural History Museum; FMNH = Field Museum of Natural History (Chicago); DMNH = Delaware Museum of Natural History; MLZ = Moore
Laboratory of Zoology (Occidental College, Los Angeles); MVZ = Museum of Vertebrate Zoology (Berkeley); NMNH = National Museum of Natural History (Smithsonian, Washington
DC). Initial Concentration in nanograms per microliter. Branch length is length of the terminal branch leading to each specimen in the maximum-likelihood tree.

These 27 individuals included 13 from the *californica* group, all from the Los Angeles basin. We sampled one specimen approximately every decade from 1880 to 2010. We call this “ancient DNA” for the purposes of consistency with the literature, although some prefer the term “historical DNA” for DNA that is not from a fresh sample, but is only hundreds as opposed to thousands of years old. One goal was to assess whether age and other factors influenced the success of data collection. Thus, our rationale for sampling all individuals from within a small area was to have one subset of our samples control for geographic variation because different lineages might have different capture efficiencies due to genomic differences (Paijmans *et al.*, 2015; Peñalba *et al.*, 2014). Geographical differences in temperature and humidity might also affect DNA degradation apart from sample age, so we also selected specimens that resided in museums in Los Angeles.

We also included nine individuals from the *woodhouseii* group and five individuals from the *sumichrasti* group, which also spanned a wide age range (1891 – 1990). Our rationale for including these samples was because, in addition to testing effects of age, we wanted to know whether the resulting data could be used to detect genetic structure at and below the species level. We also included high-quality, frozen tissue from one known hybrid individual (*californica* x *woodhouseii*) collected from a contact zone in western Nevada. For a positive control, we used tissue collected from a frozen grocery-store chicken (*Gallus gallus*).

To extract DNA from museum specimens, we cut a piece of the toe pad from each study skin using separate, sterile surgical blades in a room not used for the manipulation of PCR products. To wash away potential inhibitors of the downstream enzymatic reactions and rehydrate each toe pad, we rinsed toe pads on a Thermomixer for 5 m at room temperature in 100% ethanol, which we followed with a 5 m rinse at room temperature in 1x STE buffer (0.1 M NaCl, 10mM Trish-HCl, 0.1 mM EDTA). We minced toe pads on separate, sterile microscope slides before extracting DNA using the Qiagen DNeasy Blood & Tissue Kit protocol with modifications adapted from Mundy *et al*. (1997) and Fulton *et al*. (2012). Specifically, we extended the initial incubation step to six hours and mashed each sample with a separate, sterile mini-pestle halfway through incubation. One hour prior to the completion of incubation, we added 25 μl of 1M DTT to ensure complete digestion of toe pads. We added pre-warmed Elution Buffer (56°C) to each Qiagen column and incubated columns for five minutes at room temperature prior to elution of DNA from the membrane by centrifugation. We repeated the elution procedure into the same tube to ensure maximum recovery of DNA. For each of three chicken DNA positive controls, we extracted DNA from tissue samples using the DNeasy protocol as specified by the manufacturer in a different room from where we extracted DNA from the toe pads.

### Shearing, library preparation, and next-generation sequencing

We visualized the quality of DNA extracts through gel electrophoresis and determined DNA concentrations using a Qubit fluorometer (Life Technologies, Inc.). We also used gel images to determine the approximate DNA fragment size ranges of each sample, and we fragmented those samples whose size range was above 500 bp (the three most recent toe pad extracts and the tissue-extracted samples) on a Bioruptor (4-6 cycles on HIGH, 30 s on, 90 s off).

We prepared Illumina libraries from sheared or naturally degraded DNA samples using a KAPA library preparation kit (Kapa Biosystems), a generic SPRI substitute ((Rohland & Reich 2012); hereafter SPRI), and the with-bead (Fisher et al. 2011) library preparation method. We ligated end-repaired, adenylated DNA to Illumina TruSeq-style adapters including custom sequence tags (Faircloth & Glenn, 2012). Following a limited-cycle PCR (14 cycles) to amplify indexed libraries for enrichment, we created a library pool by combining 62.5 ng of eight indexed, individual libraries (most pools contained one chicken positive control), and we concentrated each pool to 147 ng/μl using a vacuum centrifuge. We followed an established workflow for target enrichment (Gnirke *et al.*, 2009) with modifications specified in Faircloth et al. 2012. We enriched each pool for a set of 5,060 UCEs (Faircloth *et al.*, 2012) using 5,472 synthetic RNA capture probes (MyBaits, Mycroarray, Inc.). We combined the enriched, indexed pools at equimolar ratios prior to sequencing on three runs of Illumina MiSeq PE250 sequencing (UCLA Core Genotyping Facility).

### Sequence read quality control, assembly, and UCE identification

Following sequencing, the Illumina BaseSpace platform converted BCL data to fastq format and demultiplexed each sample according to the sample-specific index we applied. We trimmed sequences for adapter contamination and low-quality bases using a parallel wrapper (https://github.com/faircloth-lab/illumiprocessor) around Trimmomatic (Bolger *et al.*, 2014). After trimming reads, we computed fastq read statistics on a per sample basis using a program (get_fasta_lengths.py) from the PHYLUCE package (https://github.com/faircloth-lab/phyluce), and we assembled contigs, *de novo*, for each sample using a parallel wrapper (assemblo_trinity.py) around Trinity (r2013-02-25) with default parameters. Following contig assembly, we conducted all data manipulations with programs available from the PHYLUCE package, unless otherwise noted. We identified contigs matching UCE loci (match_contigs_to_probes.py) in the 5k UCE locus set (https://github.com/faircloth-lab/uce-probe-sets).

### UCE processing for analysis of a concatenated data matrix

We created a “taxon set” containing the 27 species in this study and the 3 positive control samples (*Gallus gallus*), and we input this list to an additional program (get_match_counts.py) to query the database generated during UCE contig identification and create a list of UCE loci by sample. Using this list, we extracted and renamed UCE contigs from *de novo* assemblies on a sample-by-sample basis (get_fastas_from_match_counts.py), creating a monolithic fasta file for all samples during the process. We exploded the monolithic fasta file (explode_get_fastas_file.py) on a per-sample basis, and we computed UCE contig statistics from these data (get_fasta_lengths.py).

We aligned fasta sequences from the monolithic file using a parallel wrapper (seqcap_align_2.py) around MAFFT (Katoh *et al.*, 2005) that also implements a built-in trimming algorithm. This algorithm trims resulting alignments at their edges using a three-stage approach: Stage 1 removes sequence from alignment edges to ensure there are sequence data present for > 65 % of taxa that are > 75% identical over 20 bp windows; Stage 2 implements taxon-by-taxon trimming to remove alignment edges for each taxon that strongly disagree with the consensus alignment over 5 bp windows; Stage 3 re-runs the Stage 1 algorithm across the Stage 2 trimmed alignments to ensure there are sequence data present for > 65 % of taxa that are > 75% identical at alignment edges and also replaces missing data at the alignment edges with the correct encoding (“?”). We filtered the resulting alignments to create two, “*de-novo* assembled” data matrices: one that was 100% complete (no missing data for any individual) and another that was 75% complete (22 of 30 individuals must have data present in each alignment). We generated alignment statistics (get_align_summary_data.py), computed the number of informative sites (get_informative_sites.py), and concatenated all alignments in each data matrix to PHYLIP-formatted supermatrices (format_nexus_files_for_raxml.py).

### Phylogenetic analysis of concatenated UCE data

We inferred a phylogeny from the concatenated data matrix using RAxML 7.2.6 (Stamatakis, 2006) PTHREADS binary with the GTRGAMMA site rate substitution model on single, 12 CPU, 24 GB RAM nodes to conduct 20 maximum-likelihood (ML) searches for the phylogenetic tree that best fit each set of data. Following the best tree search, we generated non-parametric bootstrap replicates using the autoMRE option of RAxML, which runs bootstrapping until bootstrap replicates converge. Following the best tree and bootstrap replicate analyses, we used RAxML to reconcile the best fitting ML tree with the bootstrap replicates, and we rooted the resulting tree on the three *Gallus gallus* positive controls.

### UCE processing for SNP analysis

We selected the contigs assembled for LACMA 103135 as a reference sequence against which to align raw reads and call SNPs because this sample had the highest coverage across the largest number of UCE loci enriched. After identifying UCE loci among *de novo* assembled contigs (described above), we created a list containing the identification code for LACMA 103135, we used this list to query the database generated during UCE contig identification (get_match_counts.py), and we output a list of UCE loci for that single sample. We used this new list to extract and rename UCE contigs from *de novo* assemblies for LACMA 103135 (get_fastas_from_match_counts.py). We then created a configuration file for an automated wrapper (snps.py) around BWA (v0.7.7-r441) and PICARD (v.181) that indexes the reference contigs for alignment, uses BWA-MEM to align raw-reads to the reference data and outputs those alignments in BAM format, uses PICARD (CleanSam.jar) to check for and repair violations in the resulting BAM format, adds read group (RG) header information to each individual BAM using PICARD (AddOrReplaceReadGroups.jar), and marks duplicates in each individual BAM using PICARD (MarkDuplicates.jar). After running this program, we manually merged resulting BAMs into a single file using PICARD (MergeSamFiles.jar), and we indexed the resulting BAM using SAMTOOLS (v0.1.18). Prior to calling SNPs, we prepared the reference contigs for SNP calling by creating a sequence dictionary from them using PICARD (CreateSequenceDictionary.jar), and we indexed the reference sequence using SAMTOOLS. We used GATK (v2.7.2) to locate indel intervals (RealignerTargetCreator) in the merged BAM, realign the BAM (IndelRealigner), and call SNPs and indels (UnifiedGenotyper) in the merged BAM. We ran the resulting calls through variant filtration (VariantFiltration) to mask poorly validated SNPs (`“MQ0 >= 4 && ((MQ0 / (1.0 * DP)) > 0.1)”`), those SNPs within 5 bp of an indel, SNPs in clusters of > 10, SNP loci with QUAL under 30, and SNP loci with QD values below 2. We filtered the resulting VCF using vcftools (v0.1.12) to remove all filtered/masked SNP loci and those loci missing SNP calls for greater than 25% of all individuals. We converted the resulting VCF file to STRUCTURE and SNAPP formats using two programs from PHYLUCE (convert_vcf_to_snapp.py; convert_vcf_to_structure.py), which additionally filtered the VCF data to include only informative sites.

### Analysis of UCE SNP Data

To infer a species tree from the SNP data, we input filtered SNPs to the SNAPP (Bryant *et al.*, 2012) module for BEAST (v.2.2.1) (Bouckaert *et al.*, 2014). To be agnostic as to the species units under study, we let each individual (with its two alleles) be its own terminal tip in the species tree. We dropped the hybrid individual from the analysis (MLZ B-03). We assessed runs for convergence by visually examining posterior traces and ESS values for estimated parameters using Tracer (Rambaut et al. 2014). We determined runs had converged when ESS scores were greater than 200. To assess genetic structure by clustering analysis, we input the same set of (reformatted) data to Structure (Pritchard *et al.*, 2000), and we analyzed these data at *K* = 1–6 for 1,000,000 steps each, using a burn-in of 100,000 and correlated gene frequencies.

### Detecting contamination

Following inference of the concatenated species tree, we noticed that several individuals appeared drawn toward the outgroup/positive control (*Gallus gallus*). These samples also exhibited long terminal branches, suggesting they possessed many unique substitutions. To determine why these samples were behaving this way, we visually examined alignments using Geneious (v6.0.5) and discovered that these samples – all among the oldest – showed evidence of low-level contamination from chicken DNA toward the flanks of loci, where coverage of each sample was lowest.

We attempted to remove the contaminating sequence from several samples by aggressively trimming the edges of all contig assemblies to achieve higher coverage thresholds (minimum 3x at contig edges and 5x average). We also attempted to remove potentially contaminating reads prior to re-assembly by aligning trimmed reads to the chicken and crow genomes and filtering those reads that were more similar to chicken than to crow. This approach removed a number of putatively contaminating reads, but re-analysis of a concatenated supermatrix containing these data did not demonstrably change the phylogenetic position of the individuals and examination by eye confirmed we did not remove all contamination. We decided to use these samples to test what factors made contamination more likely. One sample (NMNH 144749) had so much contamination that we decided to drop it from the phylogenetic analyses, although we left this sample in our Structure analysis to help determine if one of the genetic clusters identified by the program reflected contaminating DNA.

### Statistical analysis

We used generalized linear models (GLM) in Stata 11 to test whether sample age had a negative effect on the number of UCE loci recovered, as well as whether sequencing effort (i.e., the total number of reads generated for a given sample) positively affected the number of UCEs enriched and the average length of those loci. We initially included the MiSeq run as an independent variable in preliminary analyses, but its effect was not significant so we dropped the variable from further analysis. We also included the interaction between sequencing effort and specimen age as an independent variable in GLMs to determine whether or not sequencing effort has a different effect depending on sample age.

We also tested for several factors that could influence contamination. We used the terminal tip length of taxa in the concatenated UCE tree as a proxy for the level of contamination, reasoning that, in the absence of contamination, individuals should cluster within their respective clades and have fairly short terminal tips – an assumption backed by previous studies of this group (Delaney *et al.*, 2008; McCormack *et al.*, 2011). Individuals with long terminal tips suggest an excess of substitutions leading to terminal taxa, which could either be due to contamination, alignment errors, or sequencing errors that make it into the consensus. Based on our observation of individual UCE alignments, most of the excess substitutions leading to long terminal tips in the concatenated UCE tree are likely due to contamination. We included the effects of sample age, sequencing effort, and initial DNA concentration as independent variables in the GLM.

## Results

### Illumina sequencing

We obtained 61 million reads across three MiSeq runs, which included the reads for the chicken positive controls. Of these 61M reads, >99% passed adapter and quality trimming, and we assembled the remaining reads into contigs for each individual (Table 1). The number of reads varied among samples by an order of magnitude, from nearly 5 million reads to fewer than 200,000 reads (average = 1.97 million reads).

### Influence of reads and age on UCE number and UCE length

We recovered as many as 4,659 (of 5,060 total = 92%) and as few as 1,515 (30%) UCE loci for the different Western Scrub-Jay historical specimens (average = 3,749 loci). We recovered 4,449 loci from the hybrid Western Scrub-Jay tissue sample and an average of 4,579 UCE loci from the three chicken positive controls. We recovered 3,423 UCEs from the oldest historical sample from 1887, and we consistently recovered around 4,000 UCE loci from specimens dating back to the late 1940s (=70 years old).

Specimen age was a predictor of the number of UCE loci we recovered (z = -9.01, p < 0.001) while sequencing effort, by itself, was not (z = -0.10, p = 0.92). However, there was an interaction between sequencing effort and age (z = 4.90, p < 0.001), in which sequencing effort had a larger effect on the number of UCEs recovered when samples were older (Fig. 1a).

**Figure 1.**
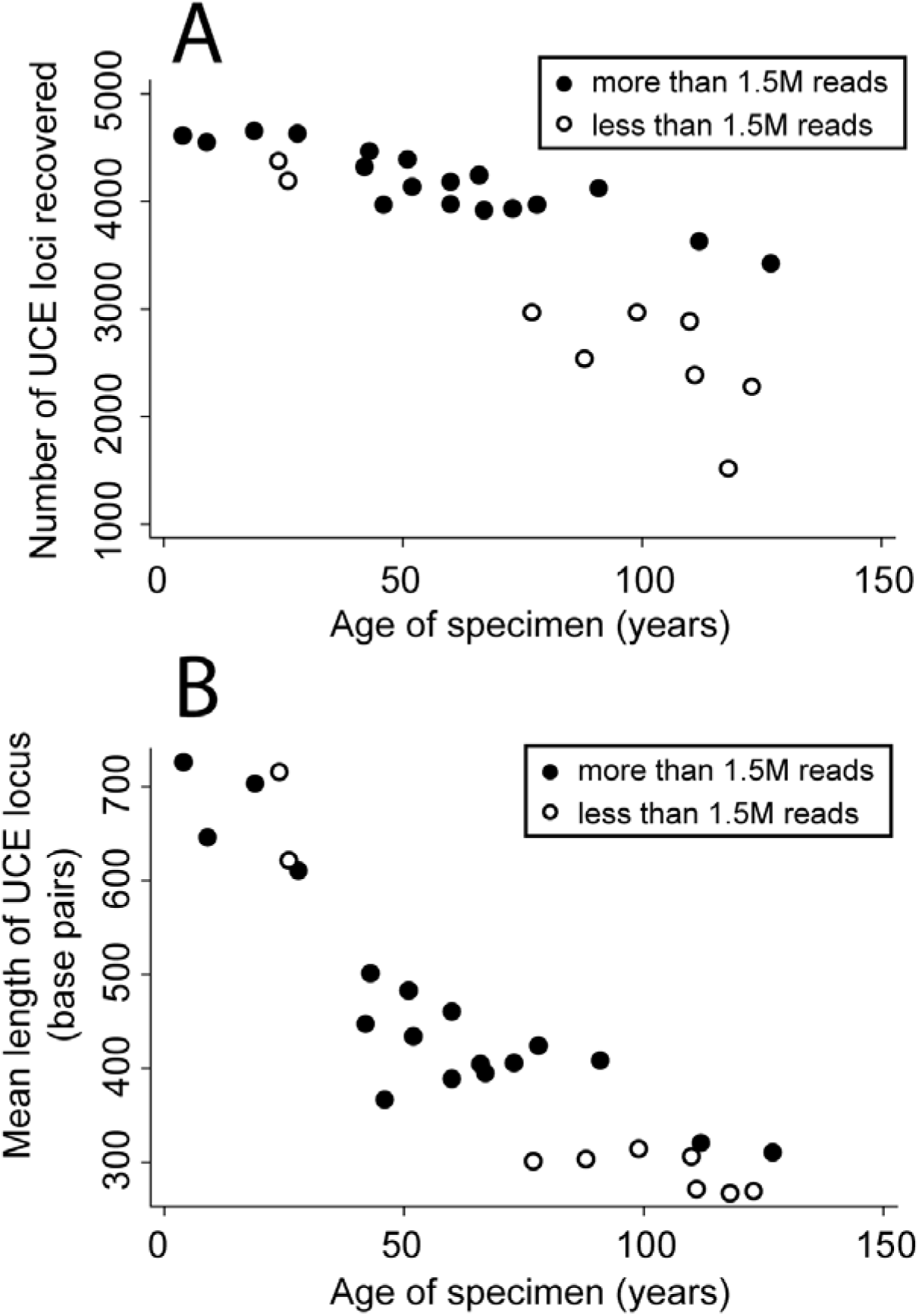
(a) The number of UCE loci recovered decreases with specimen age, but could be improved with more sequencing; and (b) the average length of UCE loci decreases with specimen age, and is less able to be improved by more sequencing.

Across all samples, the average UCE locus length was 495 bp, with a range of 267 to 867 bp. Average UCE locus length was correlated with specimen age (z = -5.78, p < 0.001), but not with sequencing effort (z = -0.60, p = 0.55; Fig 1b). In this case, there was no interaction between locus length and specimen age (z = 0.22, p = 0.83).

### Concatenation tree and species tree

Using the 75% complete data matrix of 3,770 UCE loci, we recovered a phylogeny that was broadly congruent with the known history of Western Scrub-Jays (Fig. 2). The five *sumichrasti* individuals were in a strongly supported group. Within this group, individuals from Guerrero and Oaxaca showed reciprocal monophyly. The *sumichrasti* individuals grouped together with the *woodhouseii* lineage, although *woodhouseii* and *sumichrasti* did not show reciprocal monophyly. Likewise, *californica* individuals grouped with *sumichrasti* + *woodhouseii*, again, however, without reciprocal monophyly. The known hybrid grouped with other *californica* individuals, consistent with previous results showing it contained mostly *californica* nuclear DNA (Gowen *et al.*, 2014).

**Figure 2.**
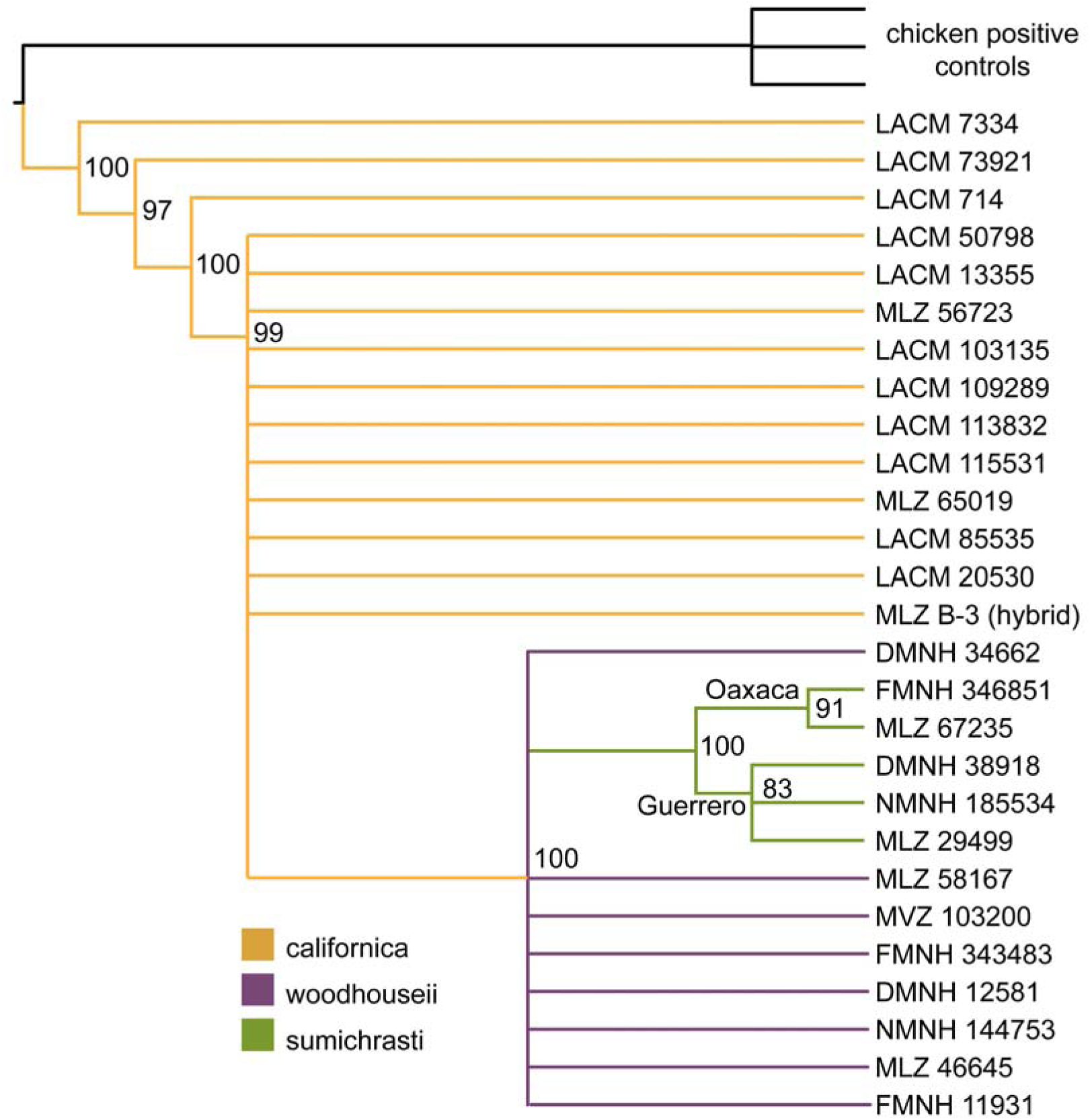
Maximum-likelihood phylogeny of Western Scrub-Jays (*Aphelocoma californica*) based on 3,770 concatenated UCEs. Nodes with lower than 75% bootstrap support are collapsed. Individuals from the three evolutionary lineages generally group close to one another, but individuals with contamination are drawn toward the outgroups. We did not include samples from the Island Scrub-Jay (*A. insularis*), which mtDNA data support as the sister taxon of the *californica* group (McCormack *et al.*, 2011) and which was previously elevated to species status.

As discussed in the methods, several factors suggest that the lack of reciprocal monophyly we observed in the concatenation tree was at least partially the result of contaminating chicken DNA pulling some samples towards the outgroup taxa. Using terminal tip length in the concatenated tree as a proxy for contamination, the GLM suggests that older samples (z = 4.02, p < 0.001), fewer reads (z = -3.57, p < 0.001), and lower DNA concentrations (z = -2.64, p = 0.008) are all predictors of contamination.

Following SNP calling, we identified 44,490 SNPs across all 28 Western Scrub-Jays. Following filtering of low-quality sites, we retained 19,387 SNPs across all lineages. The average number of high-quality SNPs per UCE locus was 4.7 (95 CI 0.10, min=1 max=27). After filtering SNPs for missing data, and including only those SNPs that were parsimony informative, the 75% complete data matrix included 4,430 high-quality, informative SNPs.

SNAPP drops loci for which there are no data for a given terminal tip (each individual in this case) – an even more conservative threshold for missing data than that which we applied above to arrive at 4,430 SNPs. This resulted in 1,388 SNPs contributing to the SNAPP analysis. We ran the SNAPP tree for 10,000,000 generations. The analysis showed signs of convergence with a stable posterior likelihood and ESS values greater than 200, except for six of the 50 theta parameters. In contrast to the topology we inferred from concatenated data, the SNAPP species tree showed strong support for monophyly of the *californica*, *woodhouseii*, and *sumichrasti* lineages (Fig. 3). Like the concatenation tree, individuals from Oaxaca and Guerrero were also placed into monophyletic groups with strong support. Within the *woodhouseii* group, the SNAPP tree showed a fine-scale geographic split between samples from northern and central Mexico, with the break occurring in northern San Luis Potosi.

**Figure 3.**
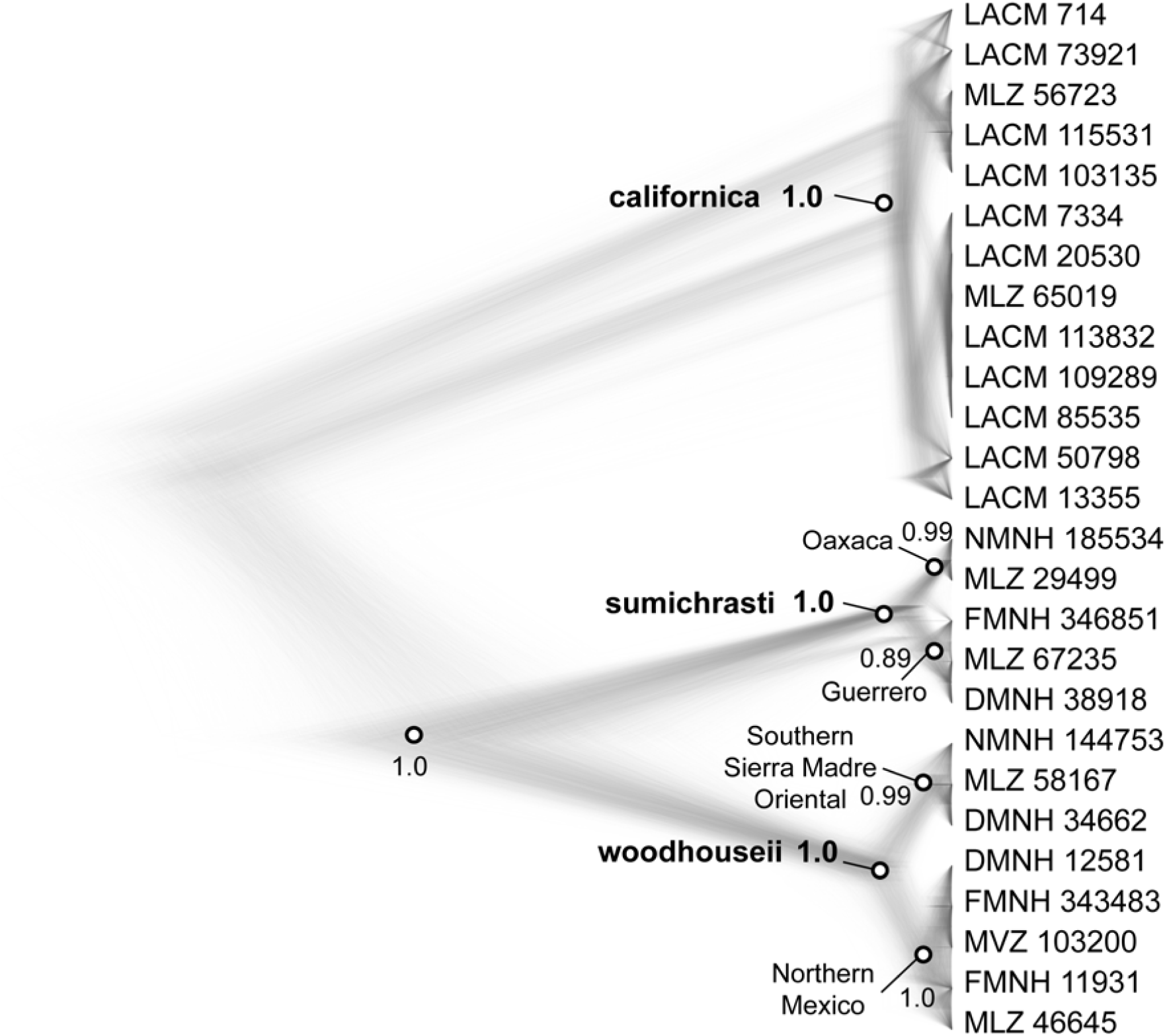
Bayesian phylogeny of Western Scrub-Jays (*Aphelocoma californica*) based on 1,388 SNPs. Unlike the maximum-likelihood tree, individuals from each evolutionary lineage cluster together in monophyletic groups. Dots represent nodes with high poster probability in the consensus tree. Important groups discussed in text are labeled. The Island Scrub-Jay (*A. insularis*) is not included in this study, but previously mtDNA results support its phylogenetic placement as the sister taxon to the *californica* group (McCormack *et al.*, 2011).

**Figure 4.**
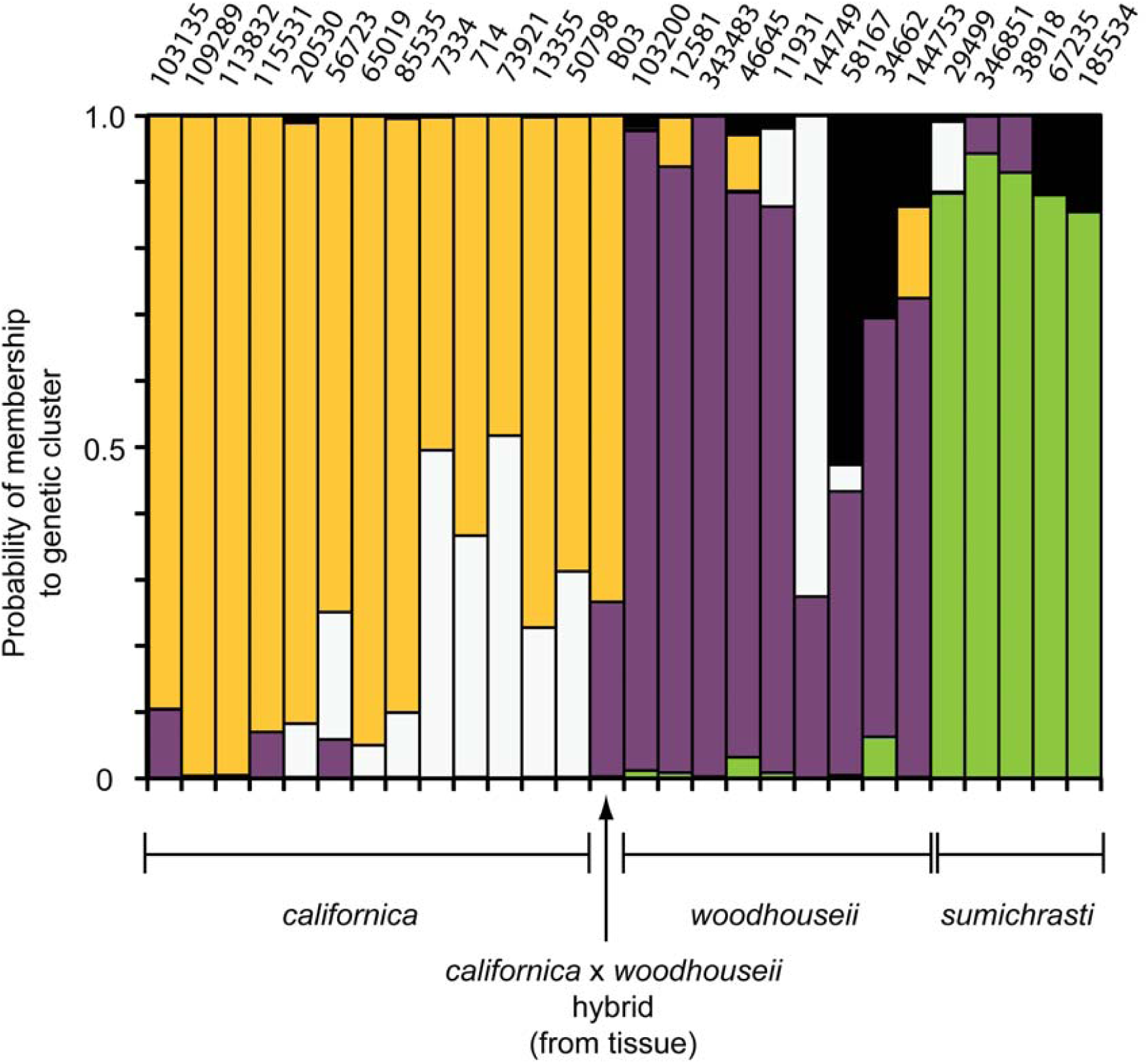
Bayesian population assignment of Western Scrub-Jays from 4,460 SNPs drawn from UCE loci. An analysis at K=5 shows three distinct clusters relating to the three main evolutionary lineages (shown in color). The fourth (white) cluster appears to reflect the level of contamination, likely from chicken DNA. The fifth cluster (black) is found in the individuals comprising the novel lineage of *woodhouseii* shown in the SNAPP tree and some assignment in two *sumichrasti* individuals.

### Bayesian clustering using SNPs culled from UCE loci

We used the 4,430 SNPs identified above in a Structure analysis. Increasing *K* values supported finer geographic structure up to *K*=5 (Average LnL over three runs at each *K*: *K*=1 -69354; *K*=2 -63703; *K*=3 -62795; *K*=4 -61003; *K*=5 -60066; *K*=6 -118881). At *K*=5, Structure assigned the five *sumichrasti* individuals, the 13 *californica* individuals, and eight of the nine *woodhouseii* individuals largely to distinct clusters. A fourth cluster appeared to reflect the amount of contaminating DNA, as it was most prevalent in the samples with long terminal tips in the concatenation tree described above. A fifth cluster was assigned with low probablity to the same *woodhouseii* individuals that formed a novel clade in the SNAPP tree. Structure assigned the hybrid individual to the *californica* cluster with an 80% probability, similar to results derived from microsatellite data (Gowen *et al.*, 2014).

## Discussion

### Age matters, especially for locus length

Ancient DNA is being increasingly studied using next-generation sequencing, but there are few studies that assess how DNA age and quality might impact different NGS approaches. A study by Paijmans et al. (2015) found that ancient DNA reacted differently than DNA from fresh samples when varying certain parameters of the sequence capture protocol. Specifically, they found that decreasing the hybridization temperature had a positive effect on capture efficiency for DNA from fresh tissue, but a negative effect on ancient DNA. This study confirms that ancient DNA might not always react as we expect with protocols developed for DNA extracted from fresh samples.

By investigating a time series of samples, we could quantify effects of sample age and sequencing effort on the number of UCE loci recovered, leading to two broad conclusions. First, we recovered fewer UCEs from older samples, but UCE recovery could be improved by more sequencing. We found that 1.5 million reads provided a rough benchmark for the sequencing effort needed to improve the recovery of UCEs. Despite some detrimental effects of age, we still recovered 2,884 loci (of 5,060 total loci) from a 110 year old sample (1904) that was subject to the shallowest sequencing (∼200,000 reads). Taken as a whole, our results suggest that sequencing effort could be reduced substantially while still recovering large quantities of data, if one is willing to accept some level of missing data. It is worth keeping in mind that more sequencing might help overcome other problems, such as the low-level contamination discussed below.

A second conclusion is that age also affects the average length of assembled UCEs. Locus length is an important metric for phylogenetic studies because longer sequences generally lead to more resolved gene trees (Castillo-Ramirez *et al.*, 2010), and, especially in the case of UCEs, there is more variability in UCE flanking regions than in UCE cores (Faircloth *et al.*, 2012). We found that average locus lengths decreased with sample age. Unlike the situation with UCE number, locus lengths were not improved dramatically by extra sequencing. For our very oldest samples (100+ years), those having more than 1.5 million reads showed modest gains in assembled locus length compared to samples having less than 1.5 million reads (300 bp versus 281 bp).

Another metric of interest is the number of enriched UCEs assembled into contigs longer than one kilobase (1 kb loci). The number of 1 kb loci we recovered from tissue samples (>1,000) reflects what we commonly see for PE250 sequencing from archived tissues. For even the most recent toe pad sample from 2010, we recovered only 322 loci greater than 1 kb. Similar to our findings with locus length, the number of 1 kb loci erodes rapidly with age. For all samples more than 45 years old, we recovered no 1 kb loci. As with locus length, increased sequencing effort did not improve the number of 1 kb UCE loci we assembled, but other solutions remain untested like using longer probes and higher probe tiling densities (e.g., Ávila-Arcos *et al.*, 2011).

One interesting question in museums is whether it is worth cutting off a portion of the specimen for cryogenic preservation to stop the steady degradation of its DNA on specimen trays. Our results suggest there may be some merit to this idea, especially for samples without archived, frozen tissue collected during the last 30 years. We observed a threshold for locus length where during the first 30 years, average length declines sharply from 867 bp to 610 bp (Fig. 1b). Then, over the next ∼100 years, locus length declines more gradually from 510 bp to just under 300 bp. Degradation will vary depending on local climate and preservation techniques, but a rough guideline is that specimens collected more recently than 1980 would be the most useful candidates for cryopreserving a small portion of the specimen (assuming no tissue was cryopreserved at the time of collection). Of course, the marginal returns in terms of slightly improved data quality should be considered relative to the cost in terms of personnel time and freezer storage space.

### Contamination

Contamination is a persistent issue in the field of ancient DNA research, even for samples younger than 100 years old. Some level of contamination is likely inevitable in ancient DNA studies (Willerslev & Cooper, 2005), although conatmination can be minimized using strict sterilization and quarantine protocols (Pääbo, 1989). Our protocols were moderately strict with respect to preventing contamination - we extracted DNA from toe pads in a different room from where we handled PCR products and performed DNA extractions from tissues. However, we included a positive control, which is not recommended (Willerslev & Cooper, 2005), and our positive control probably led to the low-level contamination we saw in some samples. These samples were marginal in terms of their initial DNA concentrations and sequencing effort. In a normal study they would probably be re-extracted, replaced with another sample, or subjected to more sequencing. For the purposes of this study, they were useful in estimating a lower bound on age, DNA quality, and sequencing effort.

For sequence capture where most read concentrate around the probe region, our study suggest that contamination will more likely manifest itself in the flanking regions of loci. Because they are a consensus assembly of many short reads, loci assembled from contaminating reads can look like chimeras. These sequences are easily detected when the contaminating DNA is phylogenetically distant from the sample DNA, as in this study. Detecting contamination would be more difficult in studies where contaminating DNA is more closely related to the focal taxa.

One might assume that initial DNA concentration is a main predictor of contamination. And while it was one predictor of contamination in this study, specimen age was the strongest predictor, followed by sequencing effort. All three variables, however, are highly correlated, and all are likely indicators of DNA quality on some level. The importance of sequencing effort is interesting because it suggests, at least for this study, that contamination occurred at a low enough level that additional sequencing could boost coverage across the flanks of UCE loci, allowing “good signal” to overwhelm background contamination.

The coalescent analysis of SNPs in the program SNAPP was better at dealing with the contamination we encountered than was the more typical UCE pipeline of calling a single allele from full-locus data and concatenating sequences for maximum likelihood analysis. For the SNAPP analysis, were were still able to place the samples with contamination into their proper evolutionary lineages, and all three lineages were strongly supported as monophyletic. The better performance of the SNP-based coalescent analysis is likely due to several factors. One, calling SNPs from UCE loci involves a hierarchical approach where the data are filtered for quality and coverage, meaning that each SNP is based on more data and more stringent parameters. It is also possible that coalescent analysis, itself, is better at dealing with the conflicting signal that results from low-level contamination, but we did not specifically evaluate this hypothesis.

### Natural history collections are genomic resources

Next-generation sequencing has the potential to transform the mission of museum collections by bringing older specimens into the age of genomics (Besnard *et al.*, 2015; Bi *et al.*, 2013; Burrell *et al.*, 2014; Nachman, 2013). To fully realize this potential, researchers need to know which NGS methods are likely to work best with DNA extracted from museum specimens and how specimen age affects data collection using different methods. For example, a recent study using RADseq to collect phylogenomic data from insect museum specimens showed that most of the RADseq data were unusuable (Tin *et al.*, 2014). While rather more success has been achieved using target enrichment approaches, studies have thus far focused on genotyping SNPs (Bi *et al.*, 2013) and not on assembling whole loci, which is likely to be more difficult with highly degraded samples.

Our results show that target enrichment of UCEs from museum specimens as old as 120 years can produce data sets including thousands of loci and thousands of informative SNPs for a range of population genetic and phylogeographic analyses. Even the oldest samples that we sequenced with relatively few reads (and which showed signs of low-level contamination) provided useful information on population assignment and evolutionary history, although other characteristics of these subpar samples would need to be interpreted cautiously, like branch lengths, substitution rates, and divergence times.

Our species tree analysis supports three distinct evolutionary lineages in the Western Scrub-Jay corresponding to the *californica*, *woodhouseii*, and *sumichrasti* groups. It had previously been unclear whether nuclear genomic data would support monophyly for these three groups, as does the mtDNA data (Delaney *et al.*, 2008; McCormack *et al.*, 2011). We did not include the Channel Island Scrub-Jay in this study, but mtDNA data support its placement as the sister taxon to the *californica* group (i.e., nested within the phylogeny). Further sampling will be needed to determine if the Guerrero and Oaxaca populations in the *sumichrasti* group (currently described as subspecies) are really as genomically distinct as the few individuals included in this study indicate. Microsatellite data, for example, suggested some gene flow between these two groups (Gowen *et al.*, 2014). The phylogenetic break in the *woodhouseii* group that we observed in the SNAPP analysis (and to a lesser extent in the Structure analysis) is novel and corresponds to a north-south split in northern Mexico, which is not reflected in current subspecies taxonomy. This pattern should be followed up with further geographic sampling.

### Recommendations

Based on results of our study, we provide recommendations for studies using sequence capture of DNA from historical museum specimens:

- More than 1.5 million reads per sample for ∼5000 targets (assuming average capture efficiency) will boost UCE recovery and, to some extent, locus length. More sequencing will also enhance good signal relative to contaminating signal in UCE flanking regions.
- Curators and collections managers should consider cryopreserving historical tissue (e.g., bird toe pads) from valuable museum specimens if the material is less than 30 years old and does not have associated frozen tissue.
- Starting DNA quality and quantity is important (also see Paijmans *et al.*, 2015). Although DNA concentration was not the best predictor of contamination, it was one predictor. Avoid using a positive control, as we did in this study, because of the high contamination risk for low-coverage samples and DNA regions (Willerslev & Cooper, 2005).
- Researchers should use strict sterilization procedures and treat DNA from museum specimens (which some prefer to call “historical DNA”) like truly ancient DNA that is thousands of years old. Low-level contamination is inevitable in some cases, but minimizing contamination during the DNA extraction and library preparation phase is the best way to avoid downstream problems. Placing an index on both sides of DNA fragments during library preparation (double-indexing) would make it easier to detect contamination that occurs after library preparation (Kircher *et al.*, 2011).
- Administrators should treat natural history collections as a genomic repository of our biodiversity, affording them the personnel needed for their protection and research use.

## Acknowledgments

We thank natural history collections and their financial supporters, collectors, collections managers, and curators for preserving a vouchered record of our biodiversity without foreknowledge of its many future uses. We thank Kimball Garrett at the Los Angeles County Museum of Natural History (LACM), Ben Marks at the Field Museum of Natural History (FMNH), Chris Milensky at the Smithsonian (NMNH), Jean Woods at the Delaware Museum of Natural History (DMNH), Carla Cicero at the Museum of Vertebrate Zoology (MVZ), James Maley at the Moore Laboratory of Zoology (MLZ), and others at those institutions who assisted with specimen loans. Uma Dandekar and Hemani Wijesuriya at the UCLA Genotyping and Sequencing Core assisted with Illumina sequencing. We thank Robert Wayne, Tom Smith, Brad Shafer, and Michael Alfaro in the UCLA Department of Ecology and Evolutionary Biology for access to a Bioruptor they purchased. Bill Mauck provided guidance on DNA extraction. Eugenia Zarza helped run the SNAPP analysis. Ed Braun provided early advice on library preparation. Brian Schmidt kindly sent photos of a Smithsonian specimen. This project was funded by NSF grant DEB-1244739 (to JEM) and the Borestone Fund of the Ridland Family (to the Moore Lab of Zoology). Computational portions of this grant were supported by DEB-1242260 (to BCF).

## Data Accessibility

DNA sequences: Genbank accessions X000000-X000000; NCBI SRA: XXX0000000

Sequence assembly, .vcf files and SNP data: DRYAD entry doi: forthcoming

R scripts and code used in the assembly: forthcoming on github

## Author Contributions

All authors designed research, WLET and BCF performed wet lab research, BCF and JEM analyzed data, all authors wrote the paper.

